# Genome analyses reveal the secondary metabolites potentially influence the geographical distribution of *Fusarium pseudograminearum* populations

**DOI:** 10.1101/2023.08.25.554839

**Authors:** Wei Li, Shulin Cao, Haiyan Sun, Xiaoyue Yang, Lei Xu, Xin Zhang, Yuanyu Deng, Igor N. Pavlov, Yulia A. Litovka, Huaigu Chen

## Abstract

Fusarium crown rot (FCR), caused by *Fusarium pseudograminearum*, significantly impacts wheat yield and quality in China’s Huanghuai region. The rapid *F. pseudograminearum* epidemic and FCR outbreak within a decade remain unexplained. In this study, two high-quality, chromosome-level genomes of *F. pseudograminearum* strains producing 3-acetyl-deoxynivalenol (3AcDON) and 15-acetyl-deoxynivalenol (15AcDON) toxins were assembled. Additionally, 38 related strains were resequenced. Genomic differences such as single nucleotide polymorphisms (SNPs), insertions/deletions (indels), and structural variations (SVs) among *F. pseudograminearum* strains were analyzed. The whole-genome SNP locus based population classification mirrored the toxin chenotype (3AcDON and 15AcDON)-based classification, indicating the presence of genes associated with the trichothecene toxin gene cluster. Further analysis of differential SNP, indel, and SV loci between the 3AcDON and 15AcDON populations revealed a predominant connection to secondary metabolite synthesis genes. Notably, the majority of the secondary metabolite biosynthesis gene cluster (SMGC) loci were located in SNP-dense genomic regions, suggesting high mutability and a possible contribution to *F. pseudograminearum* population structure and environmental adaptability. This study provides insightful perspectives on the distribution and evolution of *F. pseudograminearum*, and for forecasting the spread of wheat FCR, thereby aiding in the development of preventive measures and control strategies.

## 1. Introduction

Fusarium crown rot (FCR), caused by a complex of *Fusarium* species, is a significant disease of wheat that reduces yield and grain quality in arid and semiarid regions of the world (Kazan and Gardiner 2018; Alahmad et al. 2018). The primary pathogen of FCR is *F. pseudograminearum*, which often coexists with other *Fusarium* species, such as *F. culmorum*, *F. graminearum*, and *F. poae* (Li et al. 2012; Alahmad et al. 2018; Kazan and Gardiner 2018; Zhou et al. 2019; Deng et al. 2020; Khudhair et al. 2021). FCR caused by *F. pseudograminearum* is estimated to routinely result in a 10% yield loss in cereal crops in Australia (Murray and Brennan 2009; Alahmad et al. 2018), and up to 35% yield loss has been observed in wheat crops in the Pacific Northwest of the USA (Smiley et al. 2005). In recent years, due to changes in farming systems and global warming, FCR has continuously expanded in the Huanghuai wheat-growing region of China (Li et al. 2012; Zhou et al. 2019; Deng et al. 2020).

*Fusarium* fungi can produce a diverse range of secondary metabolites. Some of them, as pathogenic fungi, can produce different mycotoxins, such as trichothecenes (including deoxynivalenol (DON) and T-2 toxin (T-2)), zearalenone (ZEN), and fumonisin B1 (FB1) (Miller et al., 1991; Ferrigo et al., 2016; Chen et al. 2019; Li et al. 2020). These mycotoxins can help *Fusarium* infect plants, animals and even humans (Antonissen et al. 2014). Type B trichothecene mycotoxins known as DONs, further classified as 3-acetyl-deoxynivalenol (3AcDON), 15-acetyl-deoxynivalenol (15AcDON) and nivalenol (NIV), are important secondary metabolites produced by wheat pathogenic *Fusarium* species, including *F. pseudograminearum*, *F. graminearum*, and *F. asiaticum* (Ma et al. 2013; Ferrigo et al., 2016; Chen et al. 2019). The production of toxins in *Fusarium* is regulated by secondary metabolite biosynthesis gene clusters (SMGCs) (Ma et al. 2013). These clusters consist of a group of genes that work together to coordinate the production of a specific secondary metabolite (Hoogendoorn et al. 2018; Nielsen et al. 2019).

FCR caused by *F. pseudograminearum* typically occurs in wheat-growing regions that are relatively warm and dry (Kazan and Gardiner 2018). Between 2008 and 2015, 97.6% of the 297 strains of *F. pseudograminearum* isolated from Western Australia produced 3AcDON, while only 2.3% produced 15AcDON (Khudhair et al. 2019). Our previous study on the diversity of *F. pseudograminearum* strains in China indicated that strains producing 3AcDON are more prevalent in regions with higher average temperatures, such as Jiangsu Province in the south, whereas strains producing 15AcDON are more prevalent in regions with lower average temperatures, such as Shandong Province in the north (Deng et al. 2020). However, to date, there have been few studies on the impact of secondary metabolites on the environmental adaptability of *F. pseudograminearum* populations.

In this study, we selected 40 *F. pseudograminearum* strains, and divided them into 3AcDON and 15AcDON populations based on the chemotype of the mycotoxins produced by them. By performing comparative genomics analysis between these two populations, we aimed to identify genes that play a role in the evolution and environmental adaptability of different *F. pseudograminearum* populations, in addition to DON-related genes. The results of this study will help reveal the mechanisms underlying geographical distribution differences between different populations, as well as understand the field distribution and evolution of *F. pseudograminearum*.

## 2. Materials and methods

### 2.1 Fungal strains and genome sequencing

Two *F. pseudograminearum* strains CF14047 (3AcDON producer, MAT1-1) and CF16120 (15AcDON producer, MAT1-2), collected from Jiangsu and Shandong provinces, respectively, in China, were selected for de novo genome sequencing. The genomes of 38 other *F. pseudograminearum* strains with different chemotypes and mating types and the *F. asiaticum* strain CF19039 (NIV producer) were resequenced. All of the strains used in this study were single-spore isolates, and the chemotype and mating type of these strains were previously identified (Zhang et al. 2015; Deng et al. 2020). Detailed information on these strains is provided in Table S1.

Genomic DNA of these strains was extracted from mycelium prepared in liquid potato dextrose medium. The OD 260/280 ratio was detected by a Nanodrop instrument to check the purity of DNA. For de novo genome sequencing, second- and third-generation sequencing technologies were adopted. Third-generation sequencing was performed using the PacBio Sequel platform with a data depth of at least 70X. The second-generation sequencing was based on the Illumina HiSeq platform with a data depth of at least 50X. For genome resequencing, the Illumina HiSeq platform was used, and at least 3G clean data of each strain were obtained. All sequencing was performed by Genepioneer Biotechnologies Co. Ltd. (Nanjing, China).

### 2.2 Genome assembly and population variation detection

Genome assembly was performed using NextDenovo v2.0-beta.1 (https://github.com/Nextomics/NextDenovo/) based on PacBio reads (long, less accurate), and corrected using NextPolish v1.4.0 (https://github.com/Nextomics/NextPolish) (Hu et al. 2020) based on the PacBio reads and Illumina reads (short, accurate). The genome of *F. pseudograminearum* strain CS3096 (Gardiner et al. 2018) was used as a reference. Contigs obtained from the first two steps were further analyzed in Geneious Prime 2022.1.1 (Kearse et al. 2012). If necessary, the Illumina reads were mapped to the contigs with Bowtie2 (Langmead and Salzberg 2012) and SAMtools (Li et al. 2009) for confirmation. After several iterations, the genomes of CF14047 and CF16120 were finally determined. The protein-coding genes in the genome were predicted using Augustus (v3.3.3, https://github.com/Gaius-Augustus/Augustus). Homology-based gene prediction was performed using GeMoMa (v1.9) (Keilwagen et al., 2018). Multiple datasets obtained from *F. culmorum*, *F. graminearum*, *F. pseudograminearum*, and *F. venenatu* were integrated using EVidenceModeler (v1.1.1) (Haas et al., 2011). Finally, gene annotation and genome assembly quality were assessed using Benchmarking Universal Single-Copy Orthologs (BUSCO) with the Fungi_odb10 dataset (v. 4.0.6). Genome synteny and characteristic structural variations were visualized using GenomeSyn (Zhou et al. 2022).

The quality of the raw data (double-ended sequence) obtained by resequencing was assessed, and the data were filtered. Then, the clean data were aligned with the reference genome using the Burrows‒Wheeler Alignment (BWA) tool (Li and Durbin, 2009). The single nucleotide polymorphisms (SNPs) and insertion‒deletion mutations (indels) of each strain were detected and filtered based on the Genome Analysis Toolkit (GATK) (McKenna et al. 2010). The structural variations (SVs) were detected by BreakDancer (Chen et al. 2009). Based on the location of the SNPs and indels in the reference genome, annotation of these variants was performed using SnpEff (Cingolani et al. 2012). The SVs were genotyped and annotated using SVtyper v0.7.1 (Chiang et al. 2015) and SnpEff (Cingolani et al. 2012). Functional annotation of the genes containing the variation sites was carried out after filtering. By comparing genes with functional databases such as Nonredundant proteins (NR), Gene Ontology (GO), Cluster of Eukaryotic Orthologous Groups of proteins (KOG) and Kyoto Encyclopedia of Genes and Genomes (KEGG) through BLAST, functional annotations of these genes were obtained.

### 2.3 Secondary metabolite biosynthesis gene cluster annotation and analysis

The online tool antiSMASH (https://fungismash.secondarymetabolites.org/) (Blin et al. 2021) was used to search for and identify the SMGCs in the CS3096, CF14047 and CF16120 genomes. The sequences of these gene clusters were extracted from genomes using the Geneious program, and the locations were displayed on the genomes using GenomeSyn (Zhou et al. 2022). Based on the clean data obtained from the Illumina platform, the genomes of the other 38 strains were assembled de novo using SOAPdenovo2 v2.04 (Luo et al. 2012). The sequences of SMGCs in CS3096 were used as a query for a BLAST search against the genome contigs of each strain. The target gene clusters of these strains were extracted for further bioinformatics analysis.

For phylogenetic analysis, multiple alignments of SNP sequences or gene clusters were performed using the MAFFT program (https://myhits.isb-sib.ch/cgi-bin/mafft) (Katoh et al. 2005). Neighbor-joining phylogenetic trees were constructed with 1000 replicates using Molecular Evolutionary Genetics Analysis (MEGA X) software (Kumar et al. 2018). In addition, the *F. asiaticum* strain CF19039 genome was resequenced and included in the same analysis with all the sequences from *F. pseudograminearum* strains. The genome was specifically used as an outgroup in all phylogenetic analyses. The display, annotation, and management of phylogenetic trees were performed using the online Interactive Tree of Life tool (https://itol.embl.de/).

## 3. Results

### 3.1 Genome sequences of CF14047 and CF16120

The genomes of *F. pseudograminearum* strains CF14047 and CF16120 were de novo assembled using a combination of PacBio and Illumina reads. Two high quality genomes with zero-gap, telomere-to-telomere chromosomes were obtained (Table 1; Fig. 1). The nuclear genome sizes of CF14047 and CF16120 were 37.1 and 37.2 Mbp, respectively. The ends of four chromosomes in these two strains contained the telomeric sequence “GGGTTA”, and each of them was longer than that in CS3096 (Gardiner et al. 2018). A total of 11,900 protein coding genes (BUSCO completeness score, 99.7%) were predicted and functionally annotated in each genome of CF14047 and CF16120, fewer than in CS3096 (Table 1). Most of the sequences in these three genomes were syntenic, and only a few inversion blocks were found (Fig. 1). In addition to the nuclear genomes, the complete mitochondrial genomes of each strain have also been assembled. The mitochondrial genome of CF14047 is 109,151 bp in length, while that of CF16120 is 112,247 bp in length (Table 1). All of the new genome sequences of *F. pseudograminearum* have been deposited in the Genome Warehouse (GWH) of the National Genomics Data Center, Beijing Institute of Genomics, Chinese Academy of Sciences/China National Center for Bioinformation, under accession numbers GWHBOTG00000000 and GWHBOTI00000000, which are publicly accessible at https://ngdc.cncb.ac.cn/gwh/.

**Fig. 1.**
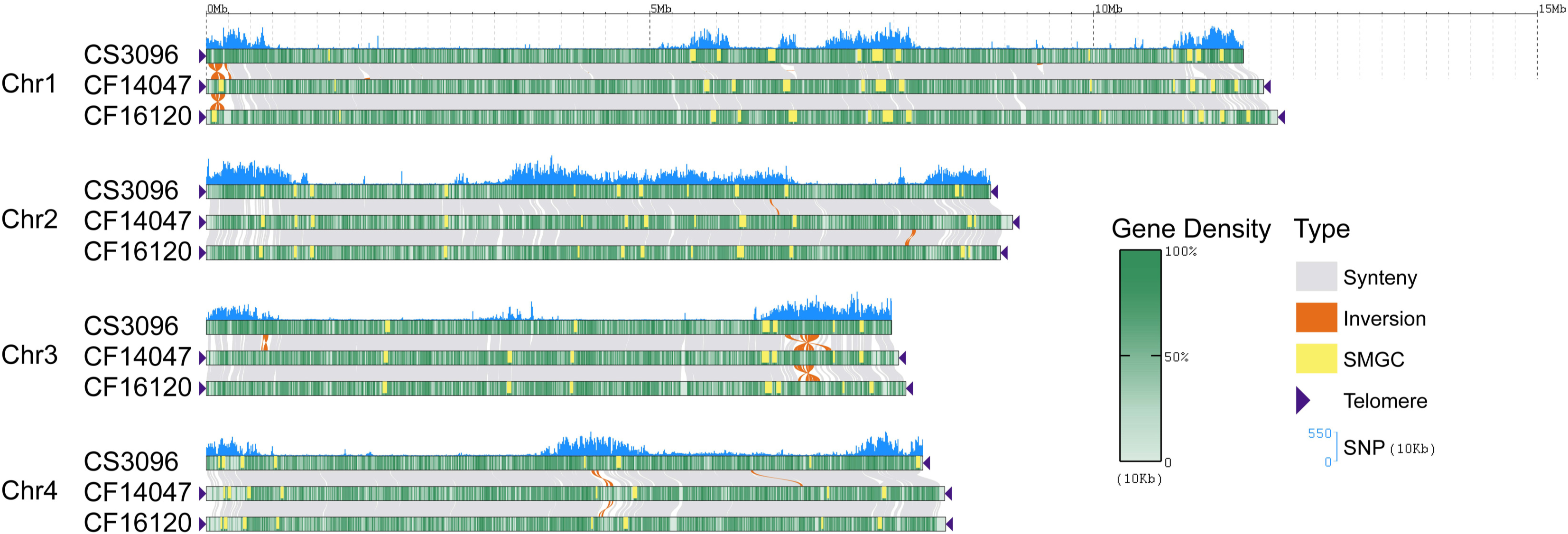
Genome synteny and characterization of structural variations among CS3096, CF14047 and CF16120.

**Table 1.**
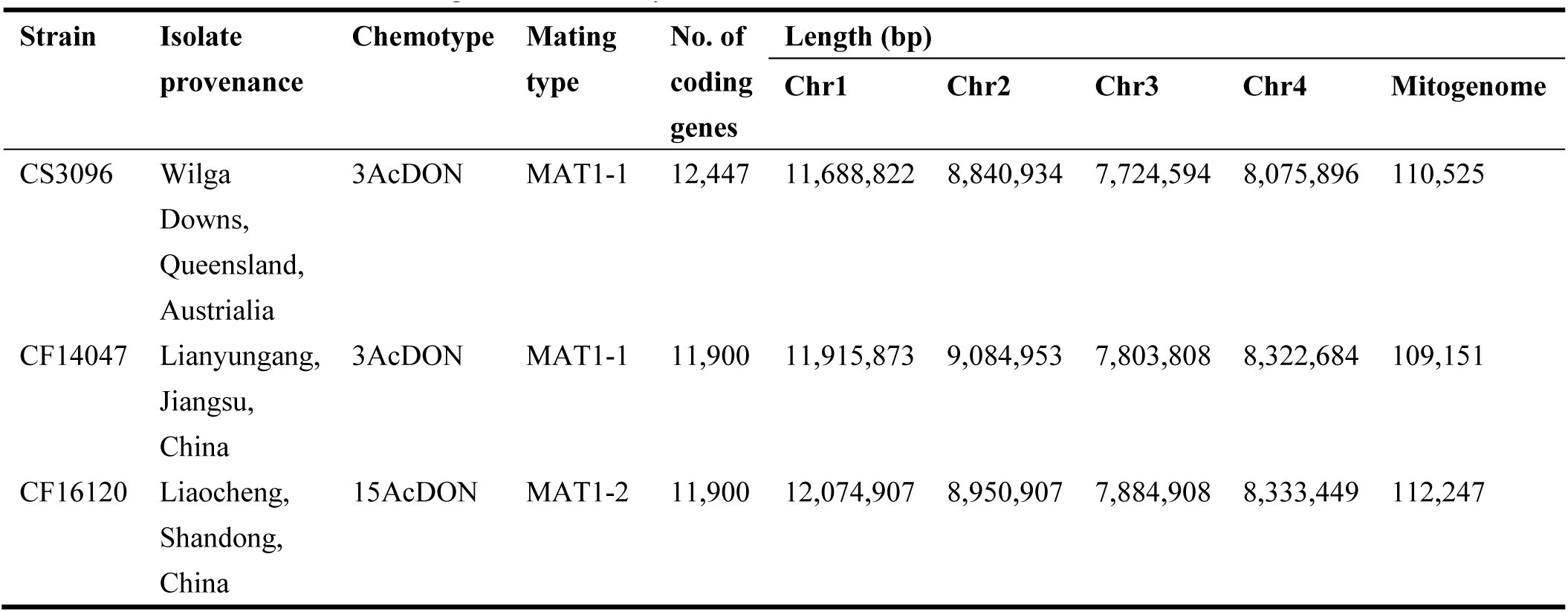
Information on the strains, genome assembly and annotation statistics.

### 3.2 Genetic variation and the annotation of different genes

In addition, 38 *F. pseudograminearum* strains were resequenced using the Illumina HiSeq platform, and an average of over 4 G clean data was obtained for each strain, with a Q30 score above 91.66%. The efficiency of sample alignment to the reference genome was 95.11%-98.03%, with an average coverage depth of 73.01X-189.17X and a genome coverage of 90.01%∼98.34% (at least one base coverage). Based on these genome data, all of the SNP, indel and SV loci in the genome and the genes containing them were searched (Table 2). In total, 296,542 SNPs were found in this study, and the density of SNPs was labeled in the genome (Fig. 1). Interestingly, the cluster analysis heatmaps based on genome-wide SNP, indel, and SV sites divide all strains into two or three groups corresponding to different chemotypes. In all three heatmaps, the 15AcDON producing strains form a single group. In the SV cluster heatmap, the 3AcDON producing strains also form a single group, and in the SNP and indel cluster heatmaps, the 3AcDON population forms two separate groups (Fig. 2).

**Fig 2.**
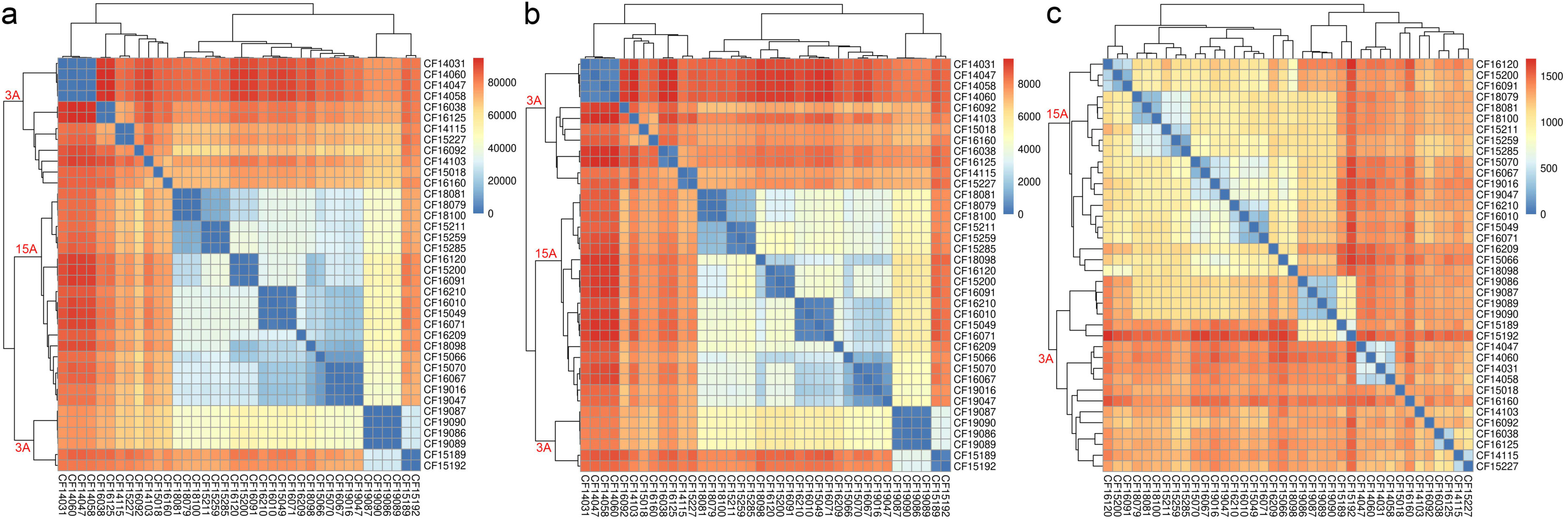
Cluster analysis heatmaps based on genome-wide SNP (a), indel (b), and SV (c) sites. 3A, 3AcDON population; 15A, 15AcDON population.

**Table 2.** The number of variations located in the nuclear genome or genes.

| **Locations** | **SNP** |  | **Indel** |  | **SV** |  |
| --- | --- | --- | --- | --- | --- | --- |
|  | **All<sup>a</sup>** | **Special<sup>b</sup>** | **All** | **Special** | **All** | **Special** |
| Genome | 296,542 | 606 | 29,565 | 54 | 3,744 | 0 |
| Gene | 10,968 | 101 | 5,544 | 25 | 850 | 0 |

a: All locations in nuclear genome or genes; b: Special locations only different between 3AcDON and 15AcDON strain populations.

Therefore, we exacted all of the SNP positions of each strain and reconstructed a phylogenetic tree based on this SNP dataset (Fig. 3). In the phylogenetic tree, all of the 15AcDON producing strains were clustered into a clade, and the rest were 3AcDON porducing strains. The strains with the same chemotype tended to cluster together; however, the clustering relationship of the strains with the same mating type was not obvious (Fig. 3). The result of phylogenetic analysis was consistent with the cluster analysis heatmaps based on differential sites, indicating that the differences in chemotypes were consistent with the differences in other sites in the genome.

**Fig 3.**
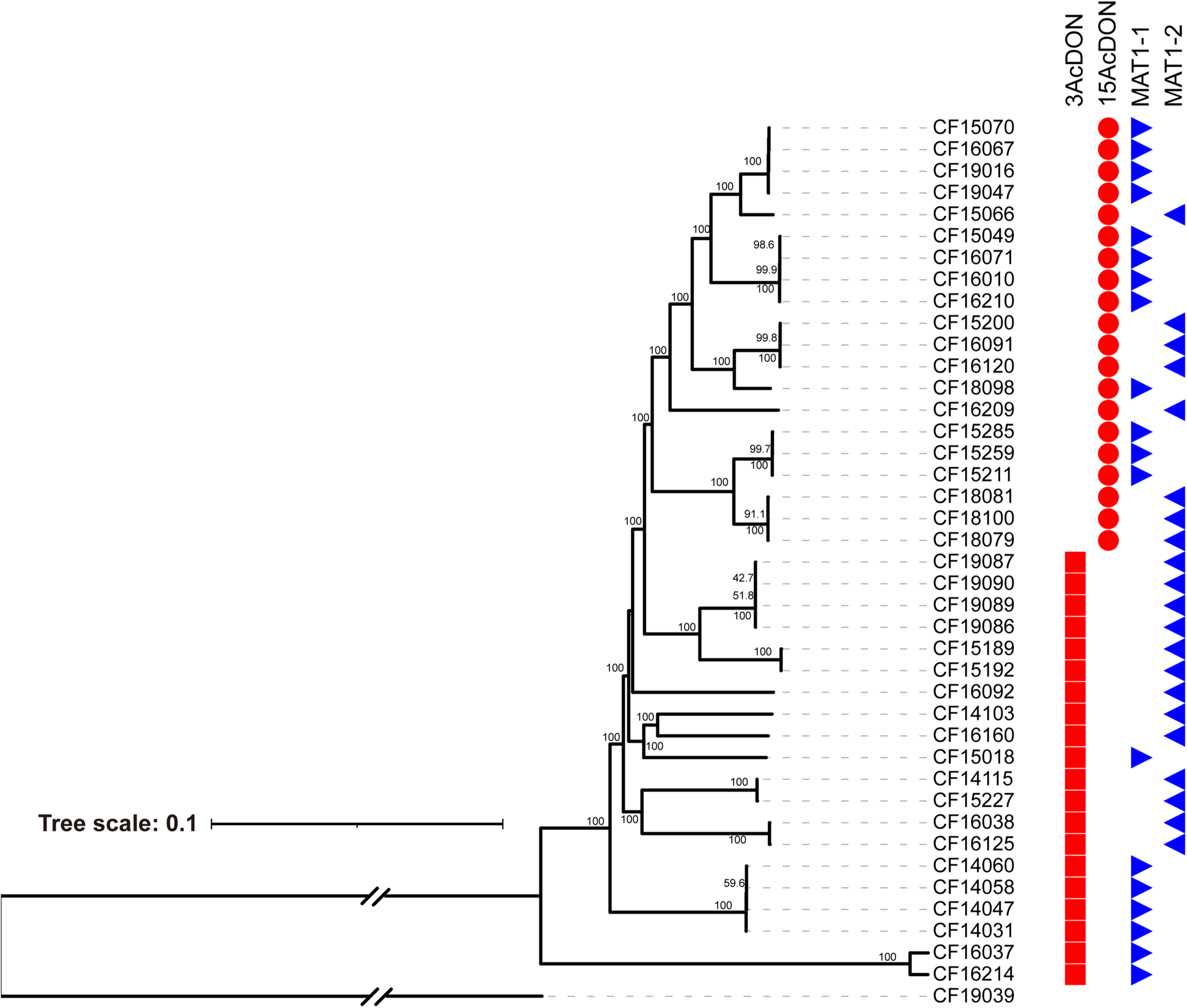
Phylogenetic tree reconstructed using the NJ method based on the SNP dataset of all *F. pseudograminearum* strains. The *F. asiaticum* strain CF19039 was used as an outgroup. The percent bootstrap values (1,000 replicates) are indicated at the nodes. The chemotype and mating type of the strains are labeled.

Based on the reference genome of CS3096, the genes containing the SNP and indel sites were annotated (Fig. 4). Among the genes associated with all SNPs and indels in the genome, the genes whose functions were related to “General function production only” were the most abundant. The genes related to “Secondary metabolites biosynthesis, transport and catabolism” were also relatively abundant, with counts of 345 (SNP) and 204 (indel), respectively (Fig. 4a, b). To analyze the genotypic differences between the 3AcDON and 15AcDON populations, we further screened the SNPs, indels and SV loci that were only different in these two populations. A total of 606 SNP sites involving 101 genes (Table S2) and 54 indel sites involving 25 genes (Table S3) were screened, and no SV sites were found to be different between the two populations (Table 2). However, among the genes associated with SNPs and indels that differed between the two populations, the genes whose functions were related to “Secondary metabolites biosynthesis, transport and catabolism” were the most important genes (Fig. 4c, d).

**Fig 4.**
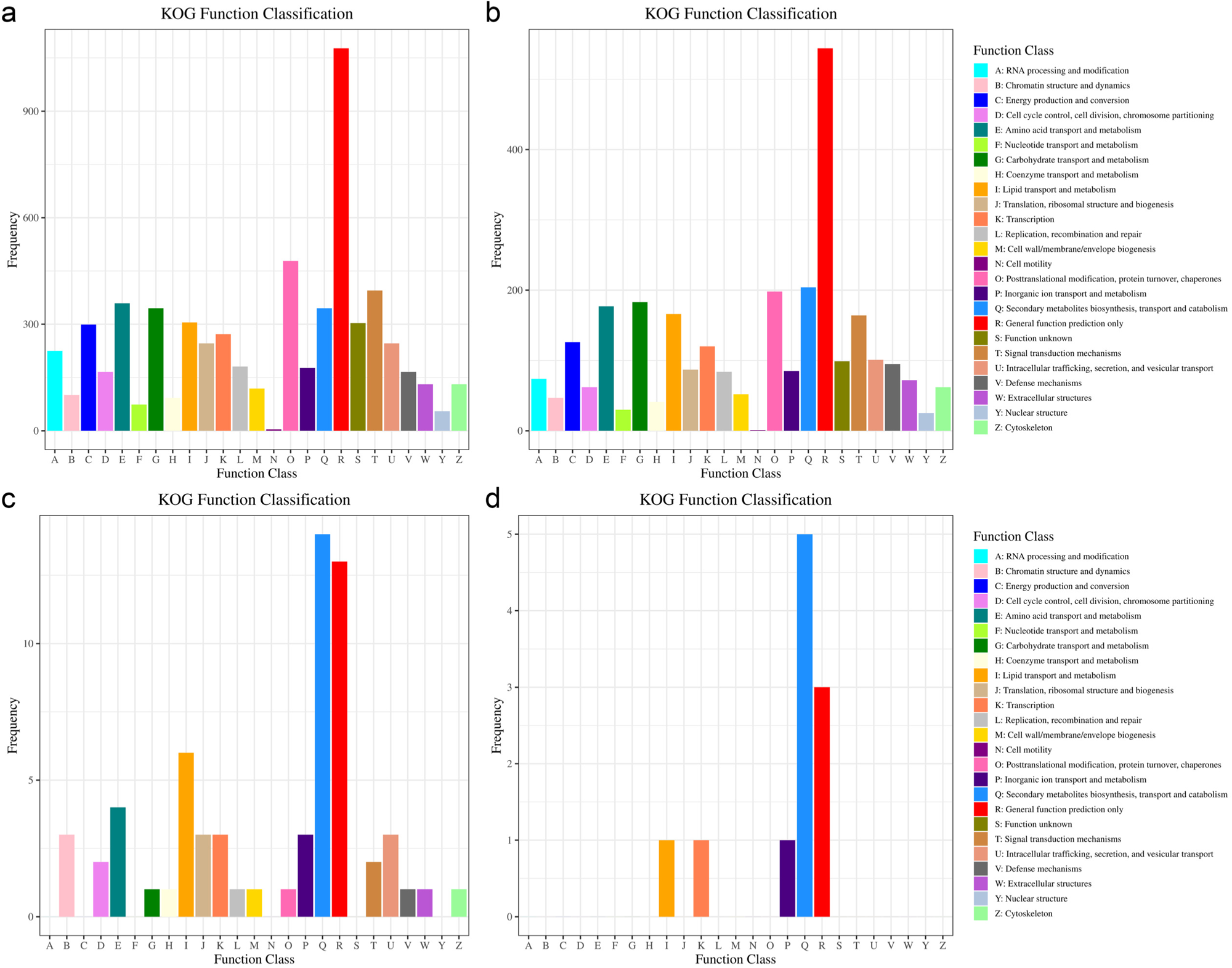
The function category of KOG annotations for the genes containing SNP and indel sites. (a) Genes containing all SNP sites. (b) Genes containing all indel sites. (c) Genes containing only SNP sites that are different between the 3AcDON and 15AcDON populations. (d) Genes containing only indel sites that are different between the 3AcDON and 15AcDON populations.

### 3.3 The difference in SMGCs between 3AcDON and 15AcDON populations

The SMGCs in the genomes of CS3096, CF14047 and CF16120 were searched using the online tool antiSMASH. There were 38 SMGCs in strain CS3096 and 40 SMGCs each in CF14047 and CF16120. In these SMGCs, nonribosomal peptide synthetases (NRPS), terpene synthases, polyketide synthases (PKS) and their hybrid compounds were commonly present in the three genomes. In addition, two other SMGCs encode cyclodipeptide synthases (CDPSs) and betalactone (Table 3). All of the SMGCs were mapped in the genomes of these three strains, and it was shown that most of the SMGCs were located in SNP-rich regions (Fig. 1). Detailed information about these SMGCs can be found in Table S4.

**Table 3.** Types and numbers of SMGCs in the three genomes of *F. pseudograminearum* strains.

| SMGS Type | *F. pseudograminearum* strains |  |  |
| --- | --- | --- | --- |
|  | CS3096 | CF14047 | CF16120 |
| NRPS | 14 | 16 | 16 |
| terpene | 9 | 9 | 9 |
| PKS | 5 | 5 | 5 |
| betalactone | 1 | 1 | 1 |
| CDPS | 1 | 1 | 1 |
| NRPS, PKS | 5 | 5 | 5 |
| NRPS, terpene | 1 | 1 | 1 |
| PKS, terpene | 2 | 2 | 2 |
| Total | 38 | 40 | 40 |

We analyzed the 25 genes containing indels that were different between the 3AcDON and 15AcDON populations. Notably, 12 genes were located in SMGCs, 8 of them in the *TRI* gene cluster (CF14047-2.8), which encodes trichothecene mycotoxins, and the other 4 genes in the CF14047-1.3, 1.6, 1.7 and 1.8 gene clusters. In contrast to the *TRI* gene cluster located on Chr2 of the *F. pseudograminearum* genome, the other four are located on Chr1 (Fig. 1; Table 4). The CF14047-1.3 and -1.6 gene clusters were NRPS or NRPS-like clusters containing one or two core genes, and the products of these gene clusters are unknown. The CF14047-1.7 and -1.8 gene clusters were NRPS and T1PKS hybrid compound clusters, each of which contained three core genes. To date, four of these gene clusters in *Fusarium* fungi have been coupled to known products, fusahexin (NRPS4) (Westphal et al. 2021), aurofusarin (PKS12) and zearalenone (PKS4+PKS13) (Nielsen et al. 2019). However, the products of the NRPS15 and PKS2 gene clusters are unknown (Table 4).

**Table 4.**
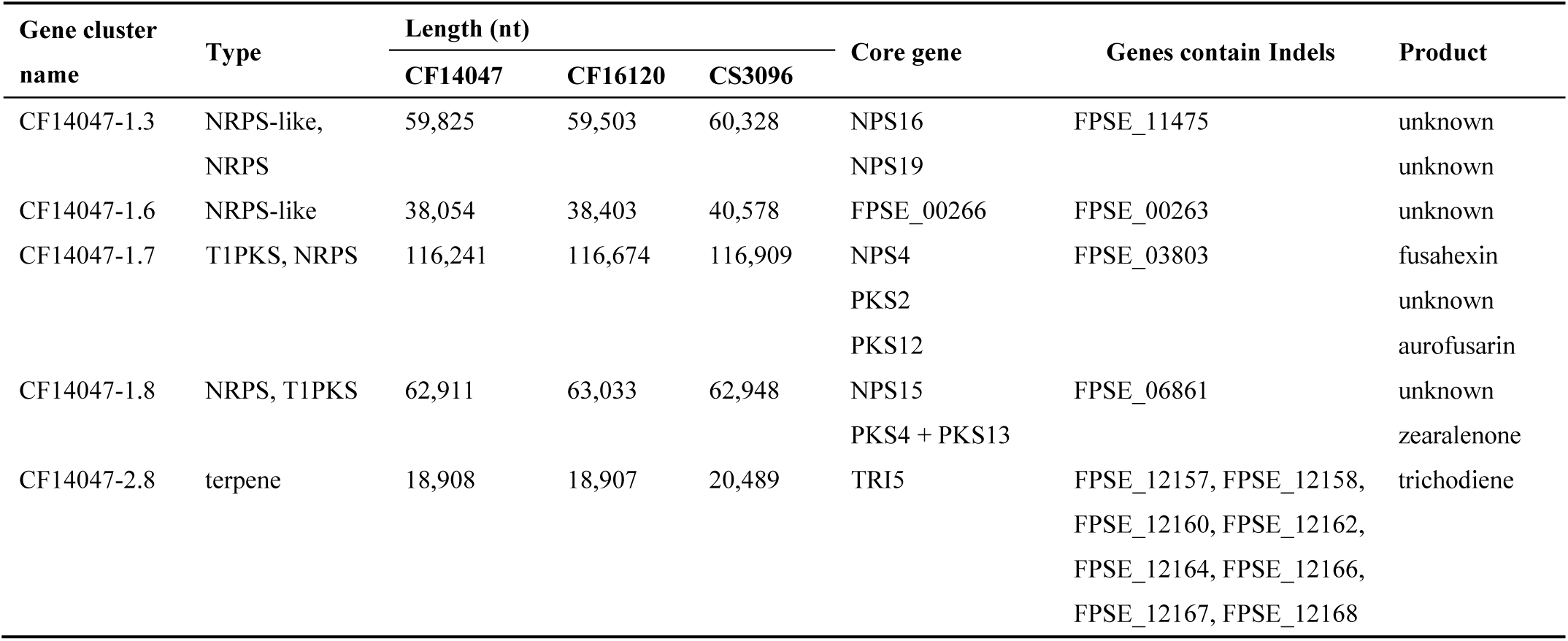
Gene clusters containing differentially expressed indel-containing genes.

We extracted the complete sequences of the 5 SMGCs (Table 4) and reconstructed phylogenetic trees based on them. In the *TRI* gene cluster tree, all strains were divided into two distinct clades corresponding to their chemotypes (Fig. 5a). In the other four trees, CF16037 and CF16214 formed a branch independent of all other strains, while the remaining strains formed roughly four larger branches. Most of these branches clustered strains from the same population (Fig. 5b-d). Strains from both populations only occurred together on branch 1 of the CF14047-1.3 tree, branches 1 and 2 of the CF14047-1.6 tree, and branch 4 of the CF14047-1.7 tree. Even on these branches, strains from the same population tended to cluster together (Fig. 5b-d).

**Fig. 5.**
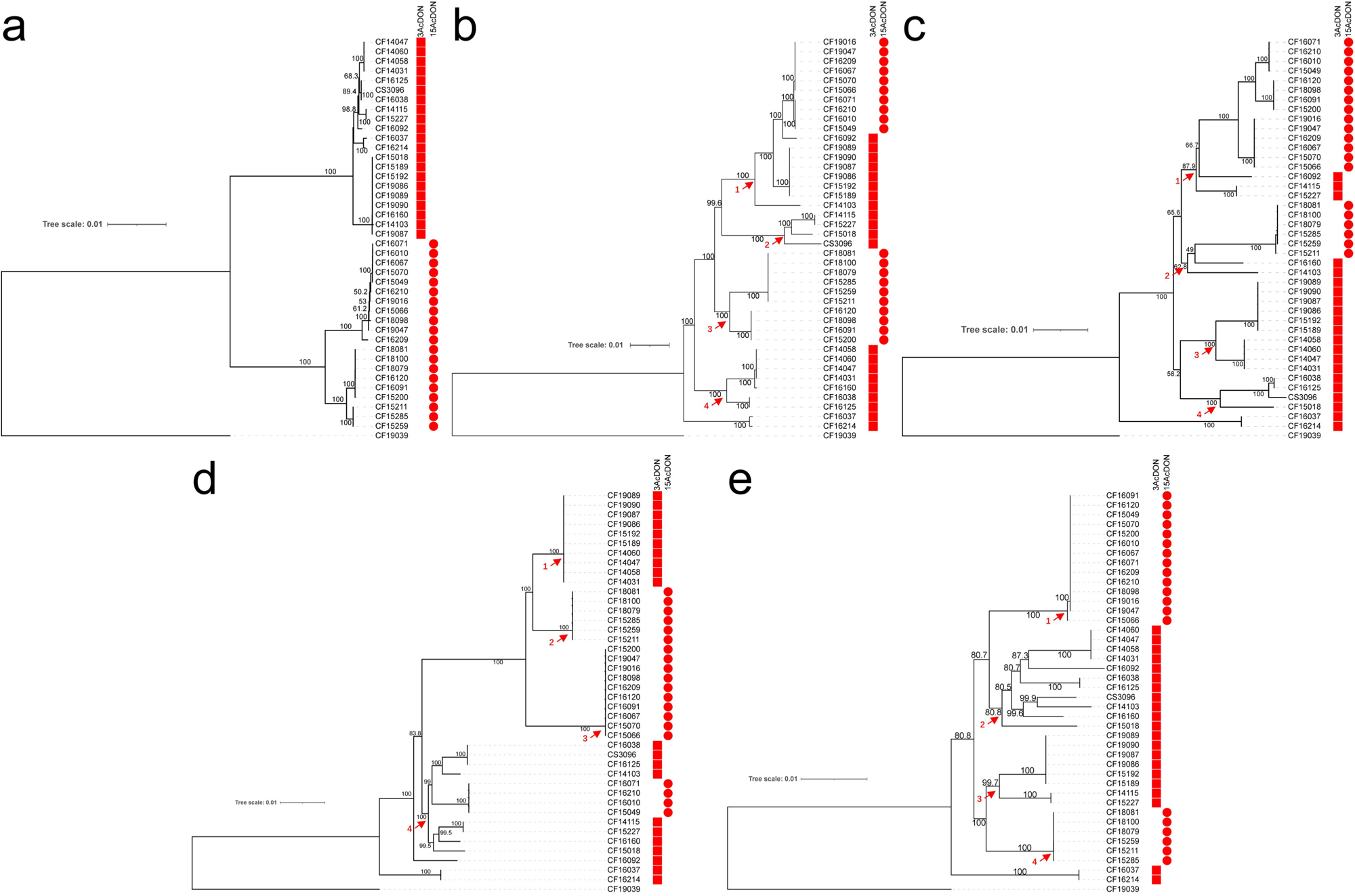
Phylogenetic analysis based on the sequences of the 5 SMGCs in the *F. pseudograminearum* genome. The *F. asiaticum* strain CF19039 was used as an outgroup. The percent bootstrap values (1,000 replicates) are indicated at the nodes. The chemotype of the strains is labeled. a, *TRI* gene cluster; b-e, CF14047-1.3, 1.6, 1.7 and 1.8 gene clusters.

## Discussion

In China, *F. pseudograminearum* was first reported as the pathogen of FCR in 2012 (Li et al. 2012). Since then, FCR caused by *F. pseudograminearum* has rapidly expanded and become the major disease in the Huanghuai wheat-growing region of China (Li et al. 2012; Zhang et al. 2015; Zhou et al. 2019; Deng et al. 2020). However, the reasons for the rapid epidemic of *F. pseudograminearum* and outbreak of FCR in China’s major wheat-producing areas within just ten years are currently unknown.

Previous studies reported that different populations of *F. graminearum* show distinct distribution patterns related to temperature, with the 15AcDON population primarily found in cooler regions of China, such as the Northeast and Huanghuaihai regions, and the 3AcDON population more prevalent in the Yangtze River and Southwest regions (Zhang et al. 2012; Shen et al. 2012). While the 3AcDON producing strains exhibits higher growth rates, spore production, and mycotoxin levels than the 15AcDON producing strains, there is no significant difference in their pathogenicity to wheat (Ward et al. 2008; Liu et al. 2017). For *F. pseudograminearum*, similar patterns have been observed in the distribution of different 3AcDON and 15AcDON populations, as temperature and humidity are likely to affect the geographic distribution of these two populations (Khudhair et al. 2019, 2021; Deng et al. 2020).

To investigate the environmental adaptation mechanism of *F. pseudograminearum*, we performed de novo genome sequencing of two strains and genome resequencing of 38 strains. To date, many *F. pseudograminearum* strains collected from Australia and China have undergone de novo genome sequencing, with some yielding high-quality, chromosome-level genomes (Moolhuijzen et al. 2013; Gardiner et al. 2018; Shan et al. 2021; Tai et al. 2022). In this study, we used a combination of PacBio and HiSeq techniques to assemble zero-gap genomes for the 3AcDON producing strain CF14047 and the 15AcDON producing strain CF16120. The ends of all four chromosomes in these strains contained the typical telomere repeat sequence “GGGTTA”, and the complete mitochondrial genome was also assembled (Fig. 1; Table 1). Only a few *F. pseudograminearum* genomes can currently be assembled with chromosome end telomere sequences. The results of this study will help further understand the genetics and biology of *F. pseudograminearum*, as well as the differences between 3AcDON and 15AcDON producing strains.

Based on the genome data obtained in this study and the reference CS3096 genome, we searched for SNP, indel, and SV loci and their associated genes and reconstructed a phylogenetic tree based on the SNP dataset (Fig. 3; Table 2). The results showed that the population classification based on the whole-genome SNP loci was consistent with the 3AcDON and 15AcDON populations, indicating the existence of genes in the genome that were linked to the toxin gene cluster. Therefore, the environmental adaptability exhibited by the 3AcDON and 15AcDON populations is not only influenced by toxin-related genes, but may also involve other genetic differences. Apart from the *TRI* gene cluster, other differences in genes between the two populations may also be essential for *F. pseudograminearum*’s environmental adaptability. The KOG annotation performed on the genes containing all SNP and indel loci in the genome, as well as the genes containing SNP and indel loci that only appear in two populations, shows that the largest number of genes are related to “Secondary metabolites biosynthesis, transport and catabolism” (Fig. 4). Secondary metabolites generally play important roles in aspects of the survival and competitiveness of fungi, such as toxin production, plant pathogenicity, and interactions with other organisms (Li et al. 2020; Kuhnert and Collemare 2022). The findings of this study indicated that genes related to secondary metabolism may be important for understanding *F. pseudograminearum*’s evolutionary adaptation to its environment.

We searched for all SMGCs in the genomes of CF14047, CF16120, and CS3096 and marked their locations in the genomes. Interestingly, most of the SMGC loci were located in SNP-enriched regions of the genome (Fig. 1). The SNP-enriched regions in fungi have evolutionary significance, as they represent areas of the genome with higher levels of genetic variation and selection, potentially leading to adaptation and diversification of fungal populations. These regions may contain genes that are important for environmental adaptation, virulence, or other traits that affect fitness (Cuomo et al. 2007).

SNPs and indel loci with differences only between 3AcDON and 15AcDON populations were associated with 101 and 25 genes, respectively (Table 2). Generally, indels introduce larger changes in the genome, which may cause frame-shift errors or site displacement, while SNPs only introduce a change in a single nucleotide and do not disrupt the reading frame. For ease of analysis, we further analyzed only the genes associated with the 25 different indel loci in this study. Among them, 12 genes were located on 5 SMGCs. The *TRI* gene cluster contained 8 different genes, which is easy to understand because the classification of 3AcDON and 15AcDON populations is based on differences in this gene cluster. The functions and evolutionary relationships of the other 4 SMGCs with the *TRI* gene cluster may be key to understanding population environmental adaptation.

The phylogenetic analysis revealed that in the CF14047-1.3, -1.6, -1.7, and -1.8 trees, the clustering of strains from the same population was not as prominent as that in the *TRI* gene cluster tree. However, on some small branches, strains from the same population still tended to cluster together (Fig. 5). We also reconstructed phylogenetic trees based on the genes containing indels, and the topology of these trees was similar to that of trees based on complete SMGC sequences (data not shown). These results indicate that, despite the CF14047-1.3, -1.6, -1.7, and -1.8 gene clusters being located on Chr1 and the *TRI* gene cluster being located on Chr2, they may have undergone synchronized evolution to some extent. These SMGCs located in highly variable regions of the genome may have been subjected to selective pressure from environmental stress during the evolutionary process, thereby playing a role in the population formation and environmental adaptation of *F. pseudograminearum*. Further research should focus on deciphering the functions and evolution of core genes in these gene clusters.

In conclusion, we have obtained high-quality, chromosome-level genomes of two *F. pseudograminearum* strains. Through comparative genomics analysis among populations, we have demonstrated the important role of secondary metabolite synthesis genes in *F. pseudograminearum* population evolution and environmental adaptation. This study is valuable for gaining insights into the genomic distinctions between 3AcDON- and 15AcDON-producing strains of *F. pseudograminearum*, as well as for understanding the field distribution and evolution patterns of these two populations.

Table S1 Information of the strains isolated from China in this study.

Table S2 Genes containing SNPs that were only different between the 3AcDON and 15AcDON populations.

Table S3 Genes containing indels that were only different between the 3AcDON and 15AcDON populations.

Table S4 Detailed information on the SMGCs in the genomes of CF14047, CF16120 and CS3096.

